# Novel genetic tools to model functional enzyme restoration in succinic semialdehyde dehydrogenase deficiency (SSADHD)

**DOI:** 10.1101/2020.09.30.321398

**Authors:** Henry Hing Cheong Lee, Phillip Pearl, Alexander Rotenberg

## Abstract

SSADHD is a rare inborn metabolic disorder caused by the functional impairment of SSADH (encoded by the aldh5a1 gene), an enzyme essential for breaking down the inhibitory neurotransmitter *γ*-aminobutyric acid (GABA). In SSADHD, pathologic accumulation of GABA results in broad spectrum encephalopathy including developmental delay, ataxia, seizures and a risk of sudden unexpected death in epilepsy (SUDEP). Proof-of-concept systemic SSADH restoration via enzyme replacement therapy (ERT) or aldh5a1 gene transfer increased survival of SSADH knockout mice, suggesting that SSADH restoration might be a viable cure for SSADHD. However, before testing SSADH restoration therapy in patients, we must consider its safety and feasibility in context of the unique SSADHD pathophysiology. Specifically, a profound use-dependent down-regulation of GABA_A_ receptors in SSADHD indicates a risk that sudden SSADH restoration might diminish GABAergic tone and provoke seizures. Such risk may be mitigated by gradual, rather than abrupt, SSADH restoration, or by restoration that is confined to critical cell types and brain regions. We therefore describe early work to construct a novel SSADHD mouse model that allows ‘on-demand’ SSADH restoration for the systematic investigation of the rate, timing and cell-specific parameters of SSADH-restoring therapies. We aim to understand the clinical readiness of specific SSADH restoration protocols on brain physiology for purposes of accelerating the bench to bed-side development of ERT or gene therapy for SSADHD patients.

## Introduction

Succinic semialdehyde dehydrogenase deficiency (SSADHD) is a rare autosomal recessive metabolic disorder (prevalence: ∼200 documented cases worldwide) caused by loss of function mutations in the aldehyde dehydrogenase 5 family member A1 (aldh5a1) gene^1,2^. Aldh5a1 encodes SSADH, which is essential for the breakdown of the inhibitory neurotransmitter *γ*-aminobutyric acid (GABA) (**Fig. 1**). In the absence of SSADH, GABA and its metabolite *γ*-hydroxybutyrate (GHB) accumulate to pathologic levels in the brain, resulting in non-progressive broad-spectrum encephalopathy^3-5^. Paradoxically, despite profound increase in extrasynaptic GABA, patients with SSADHD experience frequent seizures and significant risk of sudden unexpected death in epilepsy (SUDEP) in a hyper-GABAergic state^6^. This is likely resultant from use-dependent compensatory downregulation of GABA_A_ and GABA_B_ receptors^7,8^. To date, treatment for SSADHD is limited to symptom relief only^9-12^. A therapy that addresses with the underlying enzyme deficiency in SSADHD is absent.

**Fig. 1:**
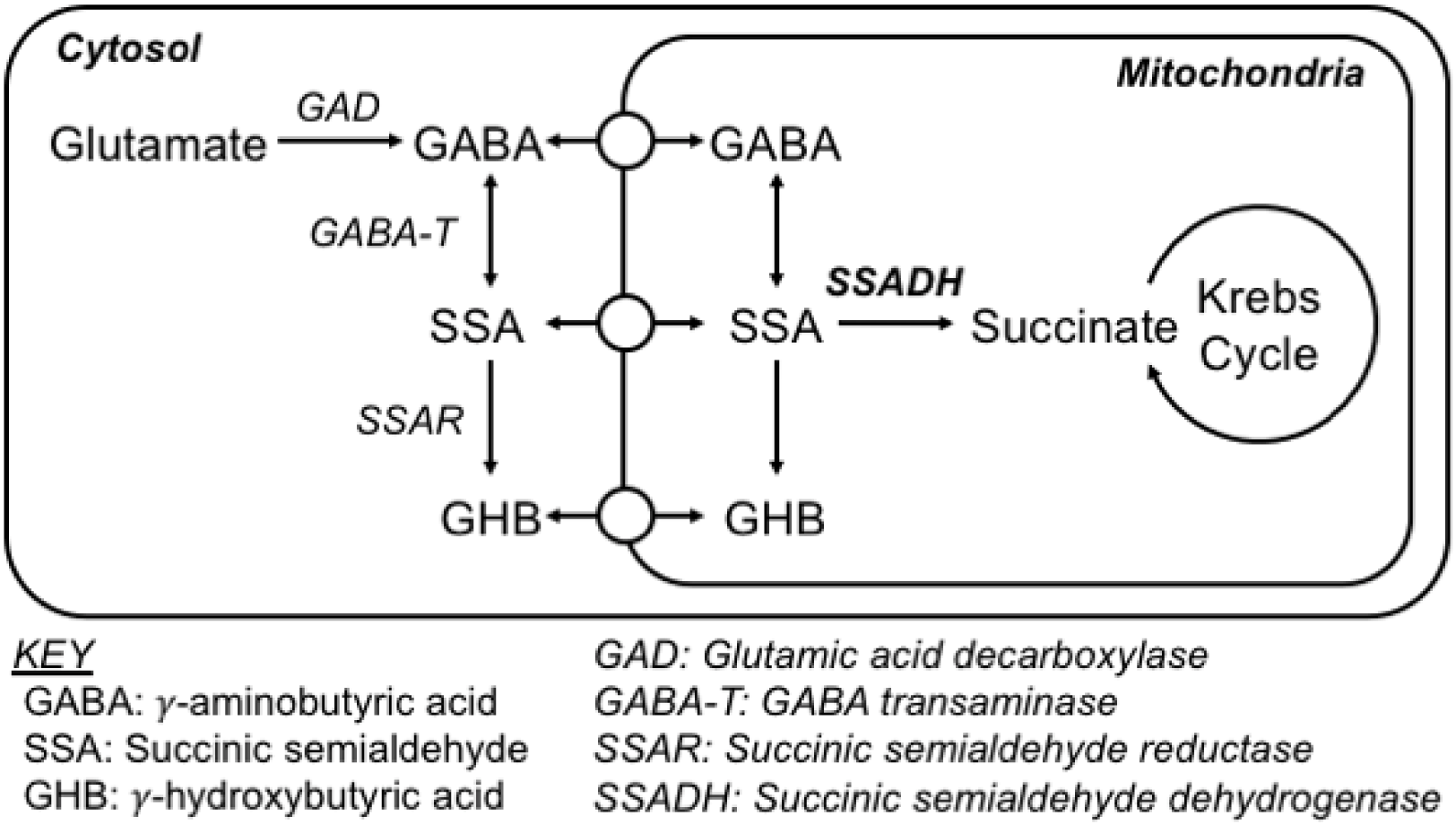
*γ*-aminobutyric acid (GABA) metabolic pathway. Cytosolic glutamate is converted by glutamic acid decarboxylase (GAD) to form GABA, which is subsequently converted by GABA transaminase (GABA-T) to succinic semialdehyde (SSA), and then by SSA reductase (SSAR) to *γ*-hydroxybutyric acid (GHB). GABA, SSA and GHB are all found in the mitochondria, where SSA is exclusively converted by SSA dehydrogenase (SSADH) into succinate, which then enters the Krebs cycle. In SSADH deficiency, GABA and GHB are accumulated in the cytosol and the mitochondria. Rational SSADH restoration strategy thus requires precise mitochondrial targeting of this key enzyme.

The aldh5a1^-/-^ mouse is the disease model of SSADHD, which mimics a severe form of the disorder^13^. Aldh5a1^-/-^ mice typically develop absence seizures at around postnatal day 15 (P15), which then progress into tonic-clonic seizures over a few days, and premature death by P25. Proof-of-concept liver-directed adenoviral aldh5a1 gene transfer^14^ or experimental enzyme replacement therapy (ERT)^15^ increases aldh5a1^-/-^ mouse survival, raising realistic prospects for clinical SSADH-restoring therapies. However, there are several major limitations pertaining to the use of this aldh5a1^-/-^ mouse tool in testing strategies that require gradual or region-specific or cell-specific SSADH restoration. First, injected enzymes elicit host immune responses that often lead to reduced therapeutic efficacy or total resistance^16,17^, limiting tests of sustained SSADH restoration. Second, functional activity of injected enzymes or viral-mediated transgene expression are uncontrollable in aldh5a1^-/-^ mice. Unmanaged SSADH restoration in aldh5a1^-/-^ mice leads to difficulty in evaluating therapeutic efficacy and dose-response relationship. Third, cell-specific SSADH restoration for therapeutic relevance is unachievable in aldh5a1^-/-^ mice without the use of viral vectors with cell-specific promoters, but available viral tools do not address various cell types relevant for SSADH expression.

Given that the full range of testing of preclinical readiness of SSADH-restoring strategies requires sustained and regulated enzyme restoration paradigms, we proposed to develop a novel SSADHD mouse model which allows conditional aldh5a1 reactivation under independent Cre regulation. In this novel mouse strain aldh5a1^lox-rtTA-STOP^, the basal activity of aldh5a1 gene is disrupted, but is reconstituted upon Cre-mediated recombination (**Fig. 2**). We will use this novel mouse genetic tool to address three key questions regarding SSADH gene therapy safety and efficacy:

**Fig. 2:**
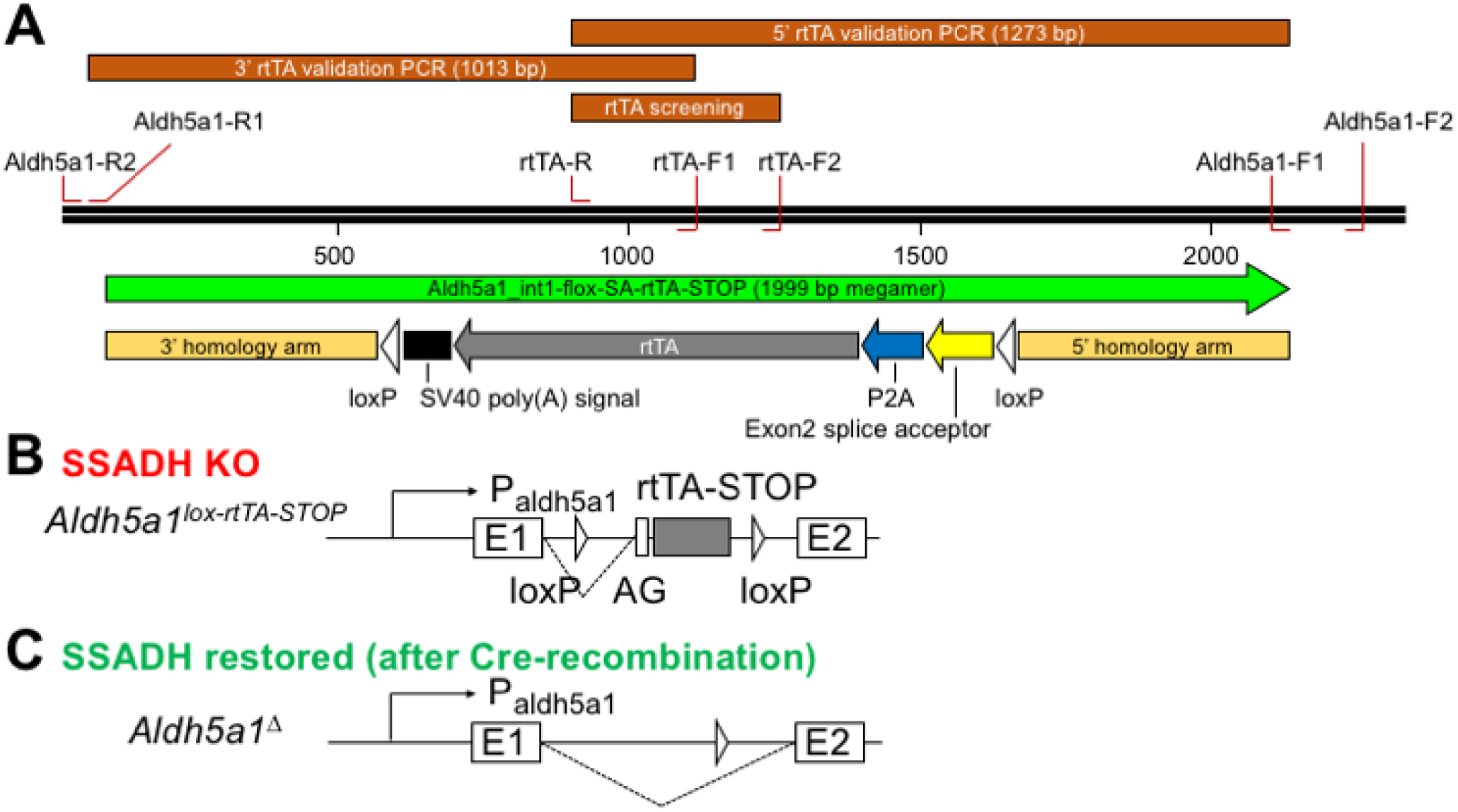
Construction and validation strategy of Aldh5a1^lox-rtTA-STOP^ mouse. **a** A single-strand DNA megamer (1999 bp) is designed for specific homology directed repair insertion into the first intron of the mouse aldh5a1gene using the Easi-CRISPR genome editing strategy. This DNA megamer contains (in the reverse strand) a cassette of splice acceptor, P2A self-cleaving peptide, reverse tetracycline transactivator (rtTA) and a SV40 poly(A) signal peptide for mRNA processing and stability, all flanked by two loxP sites. 5’ and 3’ homology arms on each end ensure specific homology directed repair insertion. Primers are designed for genomic PCR screening of candidate knock-ins. Further insert validation is performed via amplicon sequencing at both 5’ and 3’ ends. **b** At baseline, aldh5a1^lox-rtTA-STOP^ mice are SSADH-null due to early termination of the aldh5a1 gene in the presence of additional splice acceptor (AG) and downstream STOP codon. Instead, rtTA expression is driven by endogenous aldh5a1 promoter activities (for the subsequent construction of an inducible Tet-on SSADH mouse model). **c** Upon Cre-recombination, aldh5a1 is reconstituted for re-expression (aldh5a1^Δ^).

1. **How rapidly can SSADH be restored?** If SSADH restoration leads to ambient GABA reduction, then a safe rate of enzyme restoration will be determined by the maximum rate at which GABA (particularly GABA_A_) receptors are upregulated. That is, we hypothesize that abrupt SSADH restoration will correspond to abrupt GABA decline without accompanying increase in GABA receptor expression – this may lead to seizures and brain injury. In contrast, gradual SSADH replacement should enable compensatory GABA receptor upregulation, and (we predict), will be better tolerated. Using this novel mouse model, we will be the first to explicitly test the safety and efficacy of a range of rates of enzyme restoration in SSADHD (and will be the first to explicitly address *rate*, rather than *dose*, as these pertain to gene therapy for epilepsy).
2. Given tight developmental regulation of GABAergic signaling, **is SSADH restoration safe and effective across all ages?** Or is safe and effective SSADH restoration restricted to specific developmental windows? GABA circuit plasticity is heightened during early critical periods of brain development^18,19^. If successful SSADH restoration requires GABA circuit (i.e. receptor) auto regulation to accommodate a profound decline in GABA concentration, then such therapy might only be effective in younger patients. Contrarily, in older patients who lack GABA circuit plasticity, SSADH restoration might be ineffective and unsafe. This too requires explicit preclinical testing.
3. If wild-type SSADH expression is biased towards certain cell populations in the hippocampus and the cerebellum^20,21^, and global SSADH restoration risks adverse effects due to non-specific reduction in GABAergic signaling, then **is limiting SSADH restoration to relevant brain regions safer and sufficient to rescue SSADHD?** That is, we propose to test the safety and efficacy of regional and global SSADH restoration as a step toward identifying whether brain region-directed SSADH restoration (which may be safer than global SSADH restoration) is sufficient for SSADHD phenotype reversal.

## Materials & Methods

### Institutional assurance of animal and virus use

All animal treatment procedures and viral materials described in this study were covered by protocols approved by the Institutional Animal Care and Use Committee (IACUC) and the Institutional Biosafety Committee (IBC) at Boston Children’s Hospital.

### AAV injection into C57Bl/6 mice

AAV-PHP.B:CAG-GFP (2.36 x10^13^ gc/ml) was pre-packaged and obtained from the Viral Core of Boston Children’s Hospital. AAV was suspected in sterile physiological saline and was administered into C57Bl/6 mice via intraperitoneal (IP) injection at post-natal day 10 (P10). Injections were performed once or across multiple days (refer to experimental paradigms in **Fig. 5e**).

**Fig. 3:**
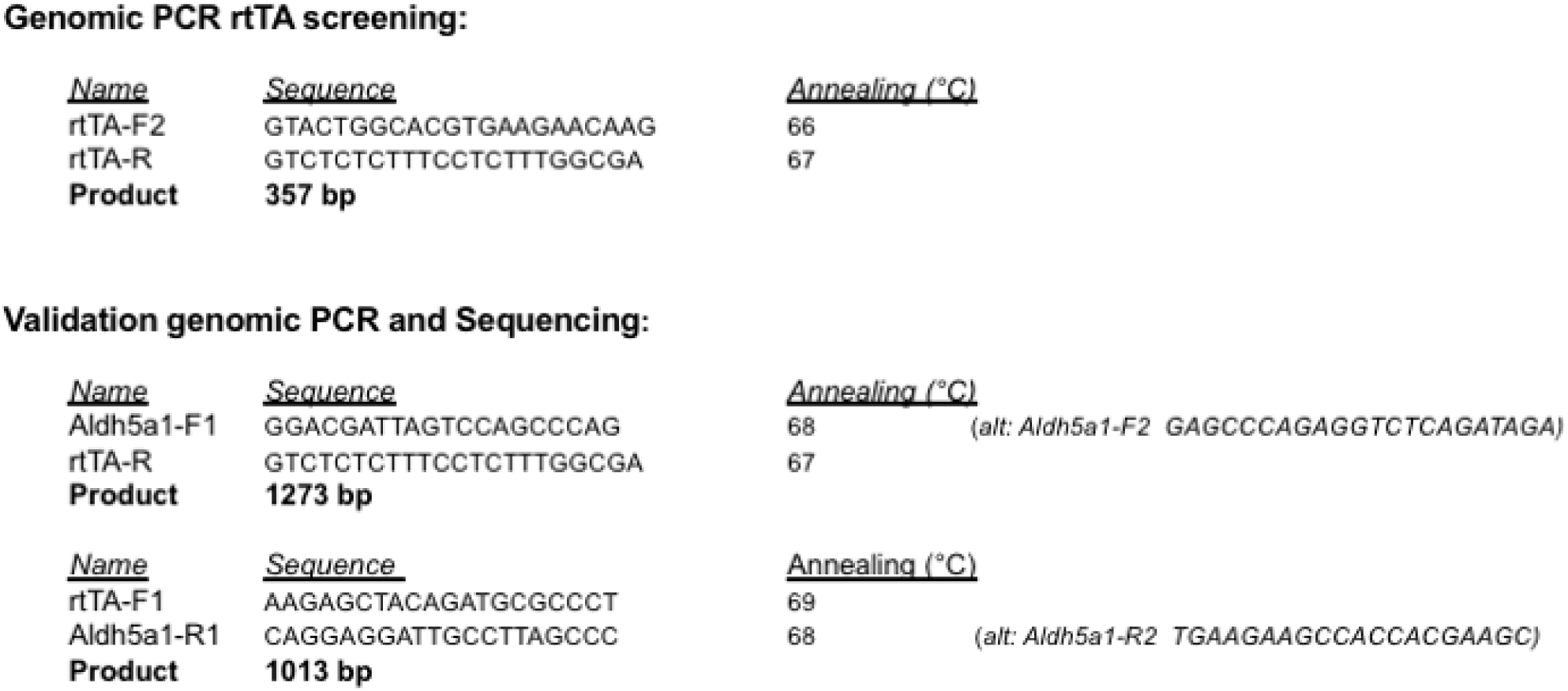
Sequence of primers for genomic PCR screening and amplicon sequencing validation for the Aldh5a1^lox-rtTA-STOP^ mouse. Names, sequences, annealing temperatures of each primer, and the PCR product sizes are listed. See **Fig. 2a** for PCR and validation strategy detail. Once this Aldh5a1^lox-rtTA-STOP^ mouse line is established, genotyping will be carried out using the genomic PCR primer pair (rtTA-F2/rtTA-R).

**Fig. 4:**
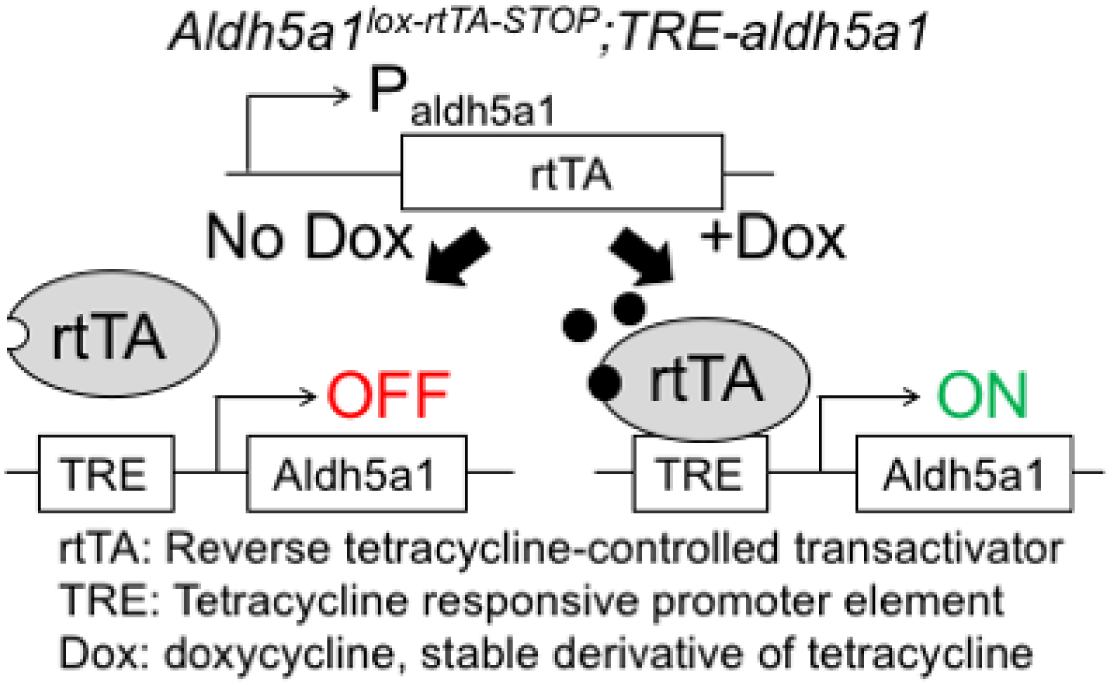
Conceptual design of the reversibly inducible SSADH mouse model. Breeding aldh5a1^lox-rtTA-STOP^ and TRE-aldh5a1 mice allows reversible expression of recombinant aldh5a1 tightly driven by a Tet-responsive element (TRE) corresponding only to the levels of doxycycline (Dox). Endogenous aldh5a1 promoter activities further limit the expressions of rtTA and recombinant aldh5a1 to relevant cell types.

**Fig. 5:**
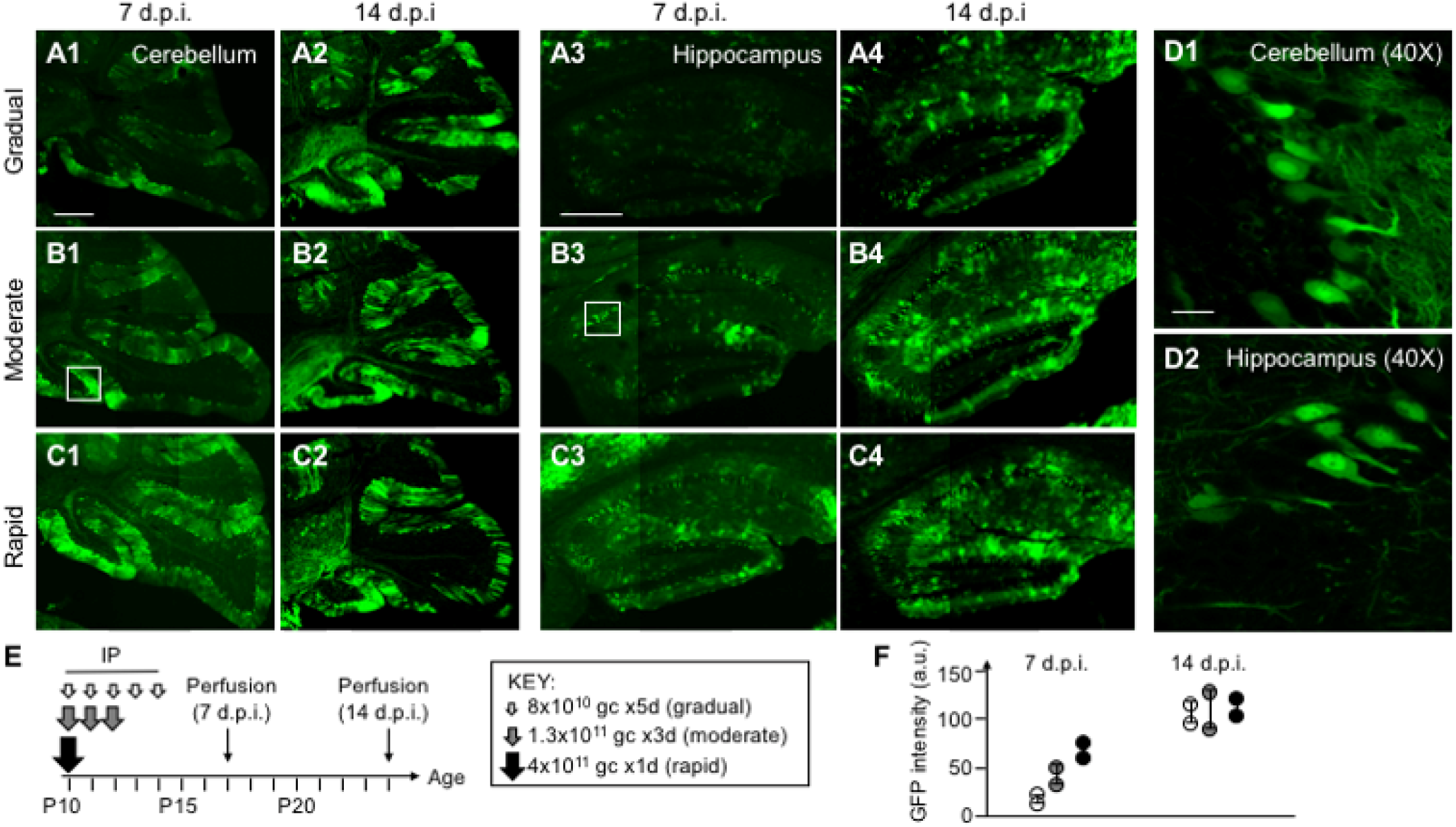
Rate-dependent GFP expression via AAV-PHP.B injected across various timespans. **a-c** Representative confocal micrographs showing the cerebellum (**a1-2, b1-2 and c1-2**) and the hippocampus (**a3-4, b3-4 and c3-4**) at 7 and 14 days post-injection (d.p.i.), in low magnification (10X), using escalating doses of AAV-PHP.B to mimic gradual (**a**), moderate (**b**) and rapid (**c**) transgene expression. **d** High magnification (40X) of individual neurons in selected brain regions in (**b1**) and (**b3**) are shown in (**d1**) and (**d2**) respectively. **e** Intraperitoneal (IP) injection paradigms across various timespans used in this study. **f** Quantification of GFP intensity in different injection paradigms and d.p.i. timepoints. Note rate-dependent expression at 7 d.p.i, which is independent from total expression at 14 d.p.i. Given the small N, no statistical analysis was performed. Scale bars: 500μm (**a1, a3**), 20μm (**d1**).

### Immunofluorescence staining

Perfusion of cortical tissue and immunostaining procedures were performed as described previously^22^. Under deep anesthesia, mice were perfused transcardially with ice-cold phosphate buffered saline (PBS) followed by 4% paraformaldehyde (PFA). Brain tissues were harvested, post-fixed in 4% PFA, and cryopreserved in Tissue-Plus OCT Compound (Fisher Healthcare, Waltham MA) for at least 24 hours before sectioning. Free-floating cryosections (coronal, 30μm, from bregma −1.23 mm through bregma −2.03 mm^23^) were obtained at −20°C, washed briefly with PBS, incubated with primary antibodies (against Calretinin, VIP or PV) overnight at 4°C, washed again, incubated with Alexa Fluor 594-conjugated secondary antibodies for 1 hour at room temperature, then mounted on glass slides. All perfusion, tissue fixation, and immunostaining procedures were carried out under the same conditions using the same batch of buffers to minimize variability between samples.

### Image Acquisition

Immunostained brain sections were identified by fluorescence imaging under low power magnification (×10 objective). Image acquisition were carried out using the FV10-ASW software (v2.1 C), with the following parameters: 20% laser output, ×1 gain control, laser intensity between 500 and 700, offset between 10% and 15% such that signals were within the linear range. Individual channels were acquired sequentially. Confocal images under low power (10X objective) and high power (40X objective) were acquired in selected brain regions.

### The novel inducible SSADH mouse model (construction work-in-progress)

We designed a two-mouse component system to allow inducible SSADH expression and depletion in a time-, rate-, and cell-specific manner. This inducible SSADH model consists of two transgenic lines: **a) the aldh5a1**^**lox-rtTA-STOP**^ **line**, in which the endogenous *aldh5a1* gene is disrupted by inserting a cassette containing a reverse tetracycline trans-activator (rtTA)/ translational STOP codon flanked by two Cre recombinase recognition (loxP) sites, and **b) the TRE-aldh5a1 line**, in which a recombined *aldh5a1* gene is inserted in a previously characterized tightly regulated (TIGRE) genomic locus for low basal expression and high inducibility^24^, to allow strict transcriptional control of aldh5a1 by a tetracycline response element (TRE) via tetracycline (or doxycycline) dosing.

The endogenous aldh5a1 gene is located in chromosome 13 (GenBank). We use a one-step mouse genome editing strategy termed Efficient Additions with Single-stranded DNA Inserts-CRISPR (Easi-CRISPR)^25^ to directly insert a lox-rtTA-STOP cassette via homology directed repair into a single-cell embryo (**Fig. 2a**). In this first mouse, aldh5a1^lox-rtTA-STOP^, endogenous SSADH expression is disrupted. To avoid alternative splicing leading to aldh5a1 gene read-through and basal SSADH expressions, we designed the lox-rtTA-STOP cassette to harbor a polypyrimidine tract directly upstream of the inserted splice acceptor site to ensure spliceosome and lariat formation^26^. The SV40 polyadenylation signal enables mRNA processing and stability. This inserted cassette ensures premature termination of endogenous aldh5a1 gene, leading to its loss of function. Instead, this mouse expresses rtTA proteins driven by the endogenous aldh5a1 promoter upon the insertion of the lox-rtTA-STOP cassette (**Fig. 2b**). Microinjection of CRISPR materials into single-cell embryos has been performed at Boston Children’s Hospital Mouse Gene Manipulation Core, and pups will be validated for gene insertion and integrity by DNA sequencing.

At basal condition, we anticipate that this mouse will phenocopy aldh5a1^-/-^, representing the severe form of the human SSADHD syndrome^13^. When injected with adeno-associated virus which encodes Cre recombinase (AAV-Cre), the rtTA cassette will be removed via Cre-lox recombination, leading to reconstituted aldh5a1 gene activities under the control of its own promoter transcriptional elements and SSADH expression restored (**Fig. 2c**). AAV-Cre will be injected at contrasting timing and dosage to test for therapeutic efficacy. The design of the aldh5a1^lox-rtTA-STOP^ mouse allows versatile approaches to further study the impacts of conditional SSADH restoration. When bred to a mouse line expressing Cre-recombinase driven by a cell-specific promoter (e.g., Gad2-IRES-Cre mouse^27^), SSADH will be restored in selective cell types. This will give insight into whether cell targeted SSADH restoration might be viable therapeutic options^28-30^. Details of genomic PCR primers and sequence validation are listed in **Fig. 3**.

The aldh5a1^lox-rtTA-STOP^ mouse will be further bred to a novel mouse line containing a tetracycline responsive element (TRE) driving a recombinant aldh5a1 gene, the TRE-aldh5a1 mouse (to be made separately) to allow reversible SSADH expression (**Fig. 4**). The recombinant gene cassette TRE-aldh5a1 will be inserted in a previously characterized tightly regulated (TIGRE) genomic locus^24^, such that SSADH expression is tightly controlled by doxycycline (dox) level. This mouse system might be particularly useful when an adaptable pace of SSADH restoration is needed over the Cre-dependent strategy (little control on AAV activities). This dox-mediated approach also allows reversible SSADH expression, so SSADH depletion can be studied systematically. Given the wide spectrum of clinical presentations among SSADHD patients traceable to their aldh5a1 mutations^31,32^, this mouse model might offer an opportunity to study individual patient’s response to SSADH replacement. Importantly, these additional experiments are unachievable using aldh5a1^-/-^ nor aldh5a1^lox-rtTA-STOP^ mouse in a Cre-dependent fashion alone (irreversible). Overall, this inducible SSADH mouse model allows controllable, reversible cell-targeted SSADH restoration which is currently unachievable using the existing tool.

## Results

### Rate-dependent transgene expression in brain via AAV-PHP.B systemic injections

A proof-of-concept study was conducted to establish experimental paradigms for various rates of transgene expression via AAV vectors. AAV-PHP.B is a recently developed capsid pseudotype with superb brain penetrance and neurotropic properties^33,34^. Using an AAV construct which expresses GFP under constitutively active promoter (AAV-PHP.B:CAG-GFP, or AAV-GFP in short), we found that transgene expression is directly proportional to the *rate* of virus vector injection. **Fig. 5** summarizes results from the pilot study where identical viral loads were delivered at once, or in 3-5 divided daily doses. We administered this AAV-GFP via IP injection in C57Bl/6 mice on postnatal day 10 (P10), and quantified AAV transduction efficiency by confocal imaging on perfused brain slices at 7- or 14-day post injection (d.p.i.). Using this injection paradigm (**Fig. 5e**), we observed widespread GFP expression in the brain, though concentrated in the hippocampus and the cerebellum, which are relevant sites of robust SSADH expression. These results are in agreement with previous AAV-PHP.B characterization suggesting robust transgene expressions throughout the brain at early time points after intravenous injections^33^. Importantly, our dosing strategies yielded differential rates of gene expression in terms of GFP intensity at 7 d.p.i. without change in cumulative GFP intensity at 14 d.p.i. (**Fig. 5f**). To our knowledge, these are the first results to indicate rate-dependent brain transgene expression after IP virus vector injection paradigm of AAV-PHP.B in developing mice.

### AAV-PHP.B transduces interneuron subtypes in the hippocampus and the cerebellum

To further characterize the cell identities of transduced cells upon AAV-PHP.B IP injections, we performed immunostaining on cryopreserved brain sections. Selected antibodies against cellular markers of different interneuron subtypes were used. Notably, we found that at 14 d.p.i., a majority of AAV-transduced GFP-expressing cells (∼80%) in the hippocampus (CA1) are calretinin positive (**Fig. 6**). In the cerebellum, however, GFP-expressing cells were VIP+ (∼60%) or PV+ (∼40%). These pilot data suggested that our experimental injection paradigms are able to transduce certain GABAergic neuron subtypes in these two investigated brain regions, using a delivery strategy (IP injection) different from those previously reported^33^. We further postulated that when this delivery paradigm is performed on the aldh5a1^lox-rtTA-STOP^ mouse using an AAV-PHP.B expressing Cre-recombinase (i.e., pAAV-CAG-Cre-WPRE-hGH, previously characterized^35^ and readily available from the Viral Core at Boston Children’s Hospital), aldh5a1 should be restored in respective GABAergic neuronal populations in corresponding brain regions. Given the relevance of predominant SSADH expression in the hippocampus and the cerebellum, and SSADH is presumably expressed in GABAergic neurons, we believe this AAV injection paradigm (combining with the aldh5a1^lox-rtTA-STOP^ mouse) will be a powerful genetic tool to model functional enzyme restoration in SSADHD.

**Fig. 6:**
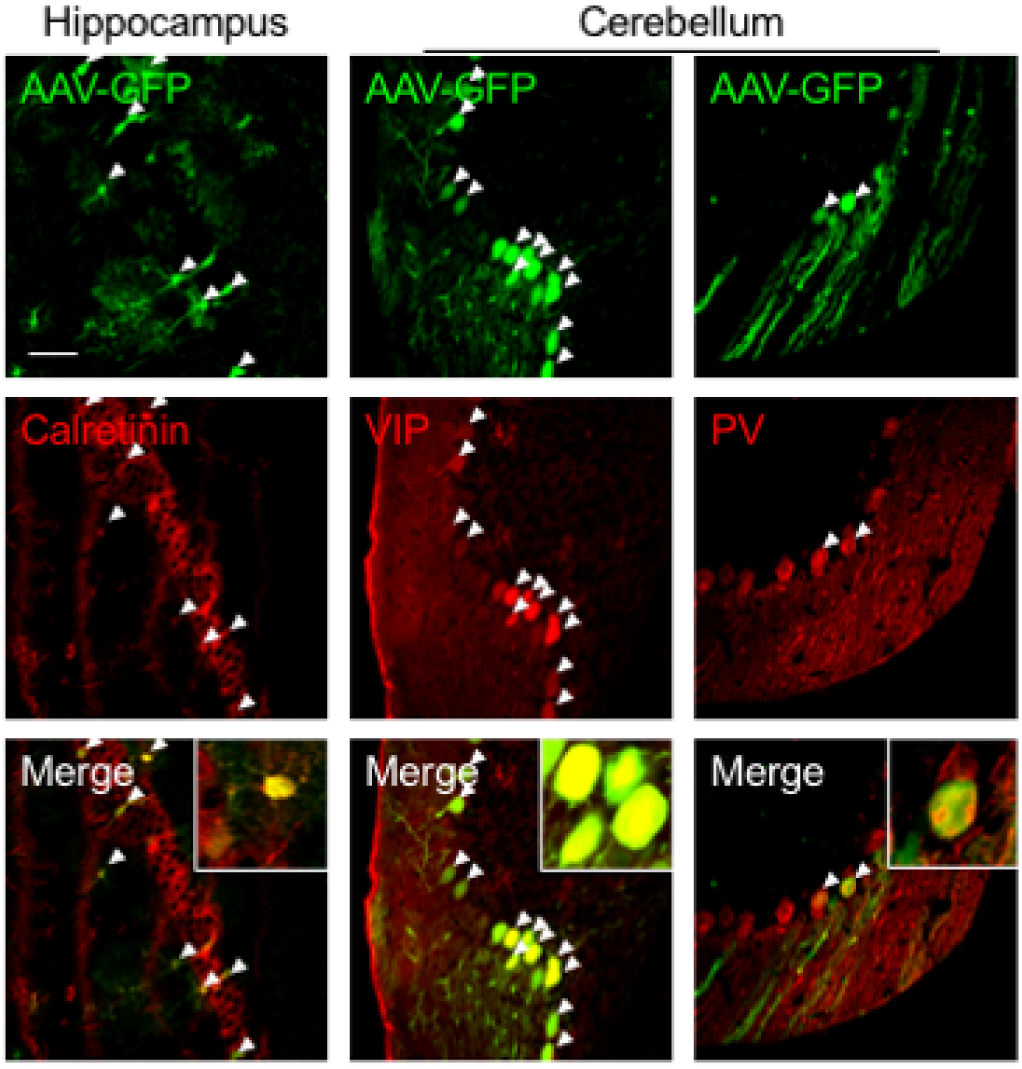
AAV-PHP.B intraperitoneally (IP) injected at P10 transduced various interneuron cell types in the brain. Representative confocal micrographs of cryopreserved brain sections showing AAV-PHP.B:CAG-GFP transduced cells (top row in green) in the hippocampus and the cerebellum. Immunostaining was performed using various interneuron cellular markers (middle row in red). Arrow heads indicate GFP-expressing cells co-immunostained by respective interneuron cellular markers (bottom row). Selected GFP+ cells are shown in high magnification in insets. VIP, vasoactive intestinal polypeptide-expressing interneurons; PV, parvalbumin interneurons. Scale bar: 50μm.

## Discussion

Our ongoing work on the aldh5a1^lox-rtTA-STOP^ mouse construction is a necessary first step to establish safety and efficacy parameters for SSADH restoration in clinical practice. We anticipate that this novel mouse tool will aid in addressing the rate, timing and cell-specific parameters for SSADH restoration. In the future, functional SSADH restoration will be further developed to achieve cell-type and sub-cellular precision. There are a few things to consider: **First**, SSADH is a mitochondrial enzyme (**Fig. 1**) with defined cell-type expression profiles^20,30^. A mitochondria-directed, cell-penetrating SSADH delivery strategy^36^ might be necessary to ensure functional SSADH restoration and to avoid off-target effects. **Second**, cell-specific SSADH delivery might be achieved via characteristic extracellular environment of relevant cell types during development^37^. For example, since the majority of PV+ cells are enwrapped by perineuronal nets (PNN) recognized by specific proteoglycan domains^38^, enzymes or viral biomolecules packaging strategy might be designed to accommodate specific extracellular interactions to target relevant cell types. **Third**, peripheral SSADH restoration might be an alternative realistic treatment option^14,15^. However, the impacts of peripheral SSADH restoration on patients’ brain physiology and long-term effectiveness must be examined in great detail. This might be addressed by the additional use of specific Cre-expressing lines with the aldh5a1^lox-rtTA-STOP^ mouse.

## Concluding remarks

We introduce a novel genetic mouse model of SSADHD, which allows ‘on-demand’ SSADH restoration and the systematic investigation of SSADH restoration parameters for preclinical readiness of ERT and gene therapy in SSADHD.

## Acknowledgments

This work is supported by a research grant from the SSADH Association (to A.R. and H.L.). We thank the Intellectual and Development Disabilities Research Center (IDDRC) at Boston Children’s Hospital (CHB IDDRC U54HD090255) for research support infrastructures, including the Viral Core for providing AAV constructs, the Animal Behavioral & Physiology Core for viral injection procedure room, and the Mouse Gene Manipulation Core for embryo microinjection with CRISPR-Cas9 materials for the aldh5a1^lox-rtTA-STOP^ mouse construction. We also thank Dr. Gerald Marsischky for consultation on the aldh5a1^lox-rtTA-STOP^ CRISPR knock-in strategy. The authors declare no conflict of interest in this study.

## References

1 Gibson, K. M. et al. Succinic semialdehyde dehydrogenase deficiency: an inborn error of gamma-aminobutyric acid metabolism. Clin Chim Acta 133, 33–42, doi:10.1016/0009-8981(83)90018-9 (1983).

2 Pearl, P. L. et al. Inherited disorders of gamma-aminobutyric acid metabolism and advances in ALDH5A1 mutation identification. Dev Med Child Neurol 57, 611–617, doi:10.1111/dmcn.12668 (2015). PMC4485983

3 Jakobs, C. et al. Urinary excretion of gamma-hydroxybutyric acid in a patient with neurological abnormalities. The probability of a new inborn error of metabolism. Clin Chim Acta 111, 169–178, doi:10.1016/0009-8981(81)90184-4 (1981).

4 Pearl, P. L., Wiwattanadittakul, N., Roullet, J. B. & Gibson, K. M. in GeneReviews((R)) (eds M. P. Adam et al.) (1993).

5 Malaspina, P. et al. Succinic semialdehyde dehydrogenase deficiency (SSADHD): Pathophysiological complexity and multifactorial trait associations in a rare monogenic disorder of GABA metabolism. Neurochem Int 99, 72–84, doi:10.1016/j.neuint.2016.06.009 (2016). PMC5028283

6 Pearl, P. L., Shukla, L., Theodore, W. H., Jakobs, C. & Michael Gibson, K. Epilepsy in succinic semialdehyde dehydrogenase deficiency, a disorder of GABA metabolism. Brain Dev 33, 796–805, doi:10.1016/j.braindev.2011.04.013 (2011). PMC4385391

7 Pearl, P. L. et al. Decreased GABA-A binding on FMZ-PET in succinic semialdehyde dehydrogenase deficiency. Neurology 73, 423–429, doi:10.1212/WNL.0b013e3181b163a5 (2009). PMC2727143

8 Parviz, M., Vogel, K., Gibson, K. M. & Pearl, P. L. Disorders of GABA metabolism: SSADH and GABA-transaminase deficiencies. J Pediatr Epilepsy 3, 217–227, doi:10.3233/PEP-14097 (2014). PMC4256671

9 Pearl, P. L. et al. Taurine trial in succinic semialdehyde dehydrogenase deficiency and elevated CNS GABA. Neurology 82, 940–944, doi:10.1212/WNL.0000000000000210 (2014). PMC3963004

10 Pearl, P. L. et al. Succinic semialdehyde dehydrogenase deficiency: lessons from mice and men. J Inherit Metab Dis 32, 343–352, doi:10.1007/s10545-009-1034-y (2009). PMC2693236

11 Vogel, K. R. et al. Thirty years beyond discovery--clinical trials in succinic semialdehyde dehydrogenase deficiency, a disorder of GABA metabolism. J Inherit Metab Dis 36, 401–410, doi:10.1007/s10545-012-9499-5 (2013). PMC4349389

12 Gropman, A. Vigabatrin and newer interventions in succinic semialdehyde dehydrogenase deficiency. Ann Neurol 54 Suppl 6, S66–72, doi:10.1002/ana.10626 (2003).

13 Hogema, B. M. et al. Pharmacologic rescue of lethal seizures in mice deficient in succinate semialdehyde dehydrogenase. Nat Genet 29, 212–216, doi:10.1038/ng727 (2001).

14 Gupta, M. et al. Liver-directed adenoviral gene transfer in murine succinate semialdehyde dehydrogenase deficiency. Mol Ther 9, 527–539, doi:10.1016/j.ymthe.2004.01.013 (2004).

15 Vogel, K. R. et al. Succinic semialdehyde dehydrogenase deficiency, a disorder of GABA metabolism: an update on pharmacological and enzyme-replacement therapeutic strategies. J Inherit Metab Dis 41, 699–708, doi:10.1007/s10545-018-0153-8 (2018). PMC6041169

16 Desai, A. K., Li, C., Rosenberg, A. S. & Kishnani, P. S. Immunological challenges and approaches to immunomodulation in Pompe disease: a literature review. Ann Transl Med 7, 285, doi:10.21037/atm.2019.05.27 (2019). PMC6642943

17 Shirley, J. L., de Jong, Y. P., Terhorst, C. & Herzog, R. W. Immune Responses to Viral Gene Therapy Vectors. Mol Ther 28, 709–722, doi:10.1016/j.ymthe.2020.01.001 (2020). PMC7054714

18 Hensch, T. K. Critical period plasticity in local cortical circuits. Nat Rev Neurosci 6, 877–888, doi:10.1038/nrn1787 (2005).

19 Le Magueresse, C. & Monyer, H. GABAergic interneurons shape the functional maturation of the cortex. Neuron 77, 388–405, doi:10.1016/j.neuron.2013.01.011 (2013).

20 Blasi, P. et al. Structure of human succinic semialdehyde dehydrogenase gene: identification of promoter region and alternatively processed isoforms. Mol Genet Metab 76, 348–362, doi:10.1016/s1096-7192(02)00105-1 (2002).

21 Zeisel, A. et al. Brain structure. Cell types in the mouse cortex and hippocampus revealed by single-cell RNA-seq. Science 347, 1138–1142, doi:10.1126/science.aaa1934 (2015).

22 Hsieh, T. H. et al. Trajectory of Parvalbumin Cell Impairment and Loss of Cortical Inhibition in Traumatic Brain Injury. Cereb Cortex 27, 5509–5524, doi:10.1093/cercor/bhw318 (2017). PMC6075565

23 Paxinos, G. & Franklin, K. B. J. Paxinos and Franklin’s the mouse brain in stereotaxic coordinates. 4th edn, (Elsevier/AP, 2013).

24 Zeng, H. et al. An inducible and reversible mouse genetic rescue system. PLoS Genet 4, e1000069, doi:10.1371/journal.pgen.1000069 (2008). PMC2346557 financial interests.

25 Quadros, R. M. et al. Easi-CRISPR: a robust method for one-step generation of mice carrying conditional and insertion alleles using long ssDNA donors and CRISPR ribonucleoproteins. Genome Biol 18, 92, doi:10.1186/s13059-017-1220-4 (2017). PMC5434640

26 Coolidge, C. J., Seely, R. J. & Patton, J. G. Functional analysis of the polypyrimidine tract in pre-mRNA splicing. Nucleic Acids Res 25, 888–896, doi:10.1093/nar/25.4.888 (1997). PMC146492

27 Taniguchi, H. et al. A resource of Cre driver lines for genetic targeting of GABAergic neurons in cerebral cortex. Neuron 71, 995–1013, doi:10.1016/j.neuron.2011.07.026 (2011). PMC3779648

28 Concolino, D., Deodato, F. & Parini, R. Enzyme replacement therapy: efficacy and limitations. Ital J Pediatr 44, 120, doi:10.1186/s13052-018-0562-1 (2018). PMC6238252

29 Toscano, M. G. et al. Physiological and tissue-specific vectors for treatment of inherited diseases. Gene Ther 18, 117–127, doi:10.1038/gt.2010.138 (2011).

30 Didiasova, M. et al. Succinic Semialdehyde Dehydrogenase Deficiency: An Update. Cells 9, doi:10.3390/cells9020477 (2020). PMC7072817

31 Akaboshi, S. et al. Mutational spectrum of the succinate semialdehyde dehydrogenase (ALDH5A1) gene and functional analysis of 27 novel disease-causing mutations in patients with SSADH deficiency. Hum Mutat 22, 442–450, doi:10.1002/humu.10288 (2003).

32 Akiyama, T. et al. SSADH deficiency possibly associated with enzyme activity-reducing SNPs. Brain Dev 38, 871–874, doi:10.1016/j.braindev.2016.03.008 (2016).

33 Deverman, B. E. et al. Cre-dependent selection yields AAV variants for widespread gene transfer to the adult brain. Nat Biotechnol 34, 204–209, doi:10.1038/nbt.3440 (2016). PMC5088052

34 Hordeaux, J. et al. The Neurotropic Properties of AAV-PHP.B Are Limited to C57BL/6J Mice. Mol Ther 26, 664–668, doi:10.1016/j.ymthe.2018.01.018 (2018). PMC5911151

35 Bei, F. et al. Restoration of Visual Function by Enhancing Conduction in Regenerated Axons. Cell 164, 219–232, doi:10.1016/j.cell.2015.11.036 (2016). PMC4863988

36 Collard, R., Majtan, T., Park, I. & Kraus, J. P. Import of TAT-Conjugated Propionyl Coenzyme A Carboxylase Using Models of Propionic Acidemia. Mol Cell Biol 38, doi:10.1128/MCB.00491-17 (2018). PMC5829486

37 Lam, D. et al. Tissue-specific extracellular matrix accelerates the formation of neural networks and communities in a neuron-glia co-culture on a multi-electrode array. Sci Rep 9, 4159, doi:10.1038/s41598-019-40128-1 (2019). PMC6411890

38 Lee, H. H. C. et al. Genetic Otx2 mis-localization delays critical period plasticity across brain regions. Mol Psychiatry 22, 680–688, doi:10.1038/mp.2017.1 (2017). PMC5400722

